# Pupation substrate and prepupal handling affect eclosion rate, timing, and adult morphology in black soldier flies (*Hermetia illucens;* Diptera: Stratiomyidae)

**DOI:** 10.1101/2025.09.15.676317

**Authors:** S.O. Durosaro, M.P. Zacarias, A. Glica, E. Atkinson, R. Rodriguez-Guevara, E. Jones, A. Gomez, M. Barrett

## Abstract

On farms, the conditions experienced by black soldier fly (BSF) prepupae may impact their development and survival, altering production goals by changing the fitness of breeding adults. Few studies have addressed the impact of environmental variables (such as substrate type or moisture content), or stressors like handling, on the development and survival of BSF as they transition from prepupae through adulthood. This study examined the effects of pupation substrate (corn cob grits, potting soil, vermiculite, wood chips, and frass), moisture content (20%, 60%, and 100%), and handling during the prepupal period (daily handling, no handling) on eclosion rate and timing, morphology (mass, head width, thorax length), and abdominal window fullness of adult BSF. Handling delayed adult emergence by over three days, reduced eclosion rates by 30.6%, reduced head width and wet mass in adult males, and reduced window fullness and head width in adult females. Frass resulted in the lowest eclosion rate (72.1 ± 2.4%) while corn cob grits (82.2 ± 2.3%) and wood chips (81.5 ± 2.2%) had the highest. Wood chips also resulted in the highest wet and dry mass, head width, and thorax length for adult males and females. Wood chips may be the best pupation substrate for BSF as it enhances body size and has a good eclosion rate. Significant prepupal handling delays adult emergence, reduces eclosion/survival rates, and reduces adult body size; however, more research is necessary to determine if the less chronic handling regimes that are likely present on farms produce similar effects.

## Introduction

Trillions of black soldier flies (BSF; *Hermetia illucens;* Diptera: Stratiomyidae) are reared each year for use as food and feed around the world; BSF are already the most abundantly farmed food or feed animal on the globe (McKay and Shah, 2025). Interest in insect farming is driven by the ability to use insects, and especially BSF larvae, as a more sustainable source of animal protein than traditional vertebrate livestock; they can also reclaim a variety of organic waste materials (Rehman *et al*., 2023; van Huis *et al*., 2013). Accordingly, BSF may be an essential part of a circular agroeconomy that minimises waste and maximises consistent, locally-available, and secure nutrition for a growing human population (Chia *et al*., 2019; Lalander et al., 2025; van Huis *et al*., 2013). However, as BSF are a relatively recent livestock animal, understanding the basic biology and welfare of this species will be key to both optimizing production and managing risk, promoting the industry’s continued growth over the next several decades (Barrett and Adcock, 2023; Barrett *et al*., 2023; Lemke *et al*., 2023).

After hatching from their eggs, BSF go through six larval instars while living in and consuming their feed (Tomberlin *et al*., 2002). The majority of BSF will be slaughtered as larvae; however, a subset of larvae must be retained, allowed to pupate and eclose, and then used for breeding as adults. These adults will lay eggs that will hatch into the next generation of larvae, continuing the colony cycle (Dortmans *et al*., 2017). To date, the majority of research into BSF production has been aimed at optimizing larval rearing. However, a facility’s overall productivity may still be impacted by conditions that affect other life stages, such as eggs, prepupae, pupae, and adults (Barrett *et al*., 2023; Lemke *et al*., 2023). As the industry has struggled with consistent adult reproduction (Lemke *et al*., 2023), determining early-life factors that may reduce viability of adult breeders, or hamper consistent reproduction, is essential to production goals.

At the end of the larval period, BSF spend 7-10 days in the ‘prepupal’ phase, where they stop feeding, become darkly sclerotised, and search for an appropriate pupation substrate in which to metamorphose. Successful pupation is essential to colony fitness and factors such as the type of pupation substrate provided and its moisture content may influence adult emergence in BSF (Dzepe *et al*., 2020; Holmes *et al*., 2013; Liu *et al*., 2023; Shumo *et al*., 2019; Singh *et al*., 2022). For example, the lack of an appropriate substrate increased the time spent in the prepupal phase, decreased the time spent during the pupal phase, and reduced adult emergence (Holmes *et al*., 2013), a pattern also observed in other species of flies (Celedonio *et al*., 1999; Sookar *et al*., 2014; Vargas *et al*., 1986). Finding appropriate pupation substrates will assist producers in optimizing adult emergence. A wide variety of substrates are reportedly used for BSF pupation on farms and in laboratories, including corn cob grits, potting soil, wood chips, vermiculite, and frass (Barrett, pers. comm); however, few have been tested for their effects on emergence and adult morphology.

Additionally, BSF pupation and eclosion rates are expected to decrease at low moisture content (e.g., <30%), as a lack of moisture can cause desiccation (Holmes *et al*., 2012; Holmes *et al*., 2013; Liu *et al*., 2023). The interactions between substrate type and moisture content have been reported to affect the time BSF spend in each developmental phase and/or mortality prior to adult emergence, as different substrates vary in their moisture retention capacities (Holmes *et al*., 2013; Liu *et al*., 2023). For example, increasing the moisture content of wood chips and vermiculite reduced BSF prepupal mortality by 95% and 88% respectively, and improved their pupation rates by 6% and 9% (Liu *et al*., 2023). However, only a few substrates have been tested for interactions with moisture content out of the many available and actively used in production facilities.

Finally, handling (e.g., shaking, sieving, or otherwise physically manipulating the animal) during the prepupal stage could result in stress that may affect subsequent pupation. Handling or other disturbances can increase octopamine (‘fight or flight’ hormones) levels in insects (Adamo and Baker, 2011; Adamo and McKee, 2017; including unpublished research in BSF: Baumann, 2019), activating increased metabolism and energy utilization as a result of adipokinetic hormone pathway stimulation (Cinel *et al*., 2020; Johnson and Barrett, 2025). Studies of chronic handling in BSF larvae have either shown more rapid eclosion (Nguyen *et al*., 2013) or no effect on eclosion and survival (Loiotine *et al*., 2024); however, larvae (unlike prepupae) have the capacity to continue eating to compensate for energy lost to the stress pathway. Chronic prepupal handling, even of a short duration, may thus have different effects on eclosion timing, survival, and subsequent adult morphology than chronic larval handling.

This study assessed the effect of five substrates (corn cob grits [CC], frass [FS], potting soil [PS], vermiculite [VM], and wood chips [WC]) at three moisture contents (20%, 60%, and 100%) and two handling treatments (daily handling of prepupae for < 20 seconds until pupation or no handling after experimental setup prior to eclosion) on developmental and morphological outcomes for BSF. These outcomes are important for understanding animal welfare (e.g., if animals are not surviving, they likely have poor welfare) and for production goals (e.g., faster colony cycles as a result of reduced development time or increased fitness as a result of larger body sizes; Addeo *et al*., 2022; Jones and Tomberlin, 2021; Knapp and Knappová, 2013). To assess how these factors influenced development, we collected data on time to eclosion and eclosion rate within 30 days. To assess how these factors influenced subsequent adult morphology, we assessed head width, thorax length, wet and dry mass, and the relative size of the fat body in the ventral abdominal window for newly-eclosed adult flies.

## Materials and Methods

### Substrates and their pretreatment

Five substrates and filter paper (#4; VWR) were used in this study: 20/40 mesh corn cob grits (CC; Blastline USA), BSF frass (FS; Fluker Farms, which was also the source of the larvae), potting soil (PS; Miracle.Gro), vermiculite (VM; MDPQT), and wood chips (WC; P.J. Murphy’s Sani-Chips). Unfortunately, we found that we could not keep the filter paper condition (meant to mimic ‘no substrate’) consistently hydrated to the correct moisture content (e.g., 20%, 60%, or 100%) like the other conditions as the single layer of filter paper in all conditions dried extremely quickly each day; therefore, data from the filter paper condition were excluded from all analyses.

We followed the protocol in Liu *et al*. (2023) to determine initial moisture content across substrates prior to re-hydration. Briefly, all the substrates were dried in a convection oven at 100°C until a constant weight was achieved for at least 4 hours (oven-dry substrate weight). As in Liu *et al*. (2023), the vermiculite was first soaked at a 1:1 vermiculite:water ratio for 24 hours before oven drying. After determining the oven-dry substrate weight, the substrates were allowed to rehydrate for 48 hr by leaving them open at room temperature and humidity (average temperature = 22.8 °C and average RH = 35% for the 48 hr); the mass of the substrate was collected a second time

(rehydrated substrate weight). Then, for each substrate, the dry-rehydrated moisture content (*R*) was calculated as:

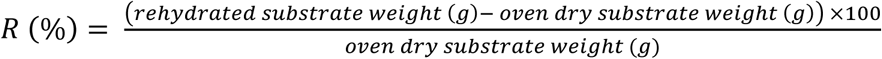

The dry-rehydrated moisture content for each substrate is shown in Supplementary Table S1. After 48 hr of rehydration, each substrate was homogenised for 5 min by mixing in a 30 L bucket (Sterilite® ClearView, USA); the container was then stored covered with an airtight lid to prevent moisture content changes prior to the experiment.

### Animal husbandry

5,000 late-stage larvae were sourced from Fluker Farms (Port Allen, Louisiana, USA). Upon arrival at the laboratory, the larvae were put into a plastic container (length 35 cm, width 24 cm, height 8.5 cm) and fed on Gainesville diet (50% wheat bran, 30% alfalfa meal, and 20% corn) at 50% moisture content until at least 50% were darkly sclerotised prepupae. Animals were then sieved from the substrate, and only mobile, darkly sclerotised prepupae were selected for the experimental setup.

### Experimental setup for substrate, moisture content, and handling

64 oz plastic cups were filled with each substrate to a depth of 8 cm (sufficient for BSF and fly pupation; Ballman *et al*., 2017; Cammack *et al*., 2010; Hou *et al*., 2006; Liu *et al*., 2023). As in Liu *et al*. (2023), the substrate’s mass was recorded, and water was added to each substrate in cups from the bottom using plastic straws attached to funnels to achieve three final moisture contents: 20%, 60% or 100% (Supplementary Table S1). The appropriate amount of water (*Y*) in mL added to each substrate to obtain the desired moisture contents (20%, 60% and 100%) was calculated using the formula:

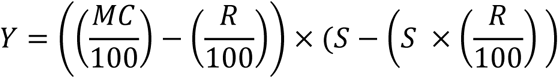

Where *MC* is the desired moisture content (%), *R* is the rehydrated dry moisture content (%), and *S* is the substrate weight in the cups. A total of six cups were made for each moisture content x substrate combination. All cups were covered with perforated lids to allow for gas exchange and placed in an incubator (Percival Scientific, I-36NL, USA) kept at room temperature. An 8.5-watt (800-lumen) LED bulb was fixed inside the incubator to supply a 14:10 L:D cycle. A temperature/relative humidity data logger (Onset HOBO data loggers S-THB-M008) was inserted in the incubator to measure the temperature and relative humidity of the incubator during the experiment (25.84 ± 0.86 °C; 71.19 ± 16.93% RH).

Twenty mobile, darkly sclerotised prepupae were weighed as a group and transferred into each cup, resulting in 1,800 prepupae in total. Each substrate x moisture combination (n = 6 cups) was then further divided into 3 handled and 3 non-handled replicates. Every 24 hours for 30 days, all replicates were removed from the incubator. Handled cups were then sieved for approximately 20 seconds to separate the prepupae from the substrate. Substrate was then returned to the cup and the number of prepupae and pupae were counted. Any pupae found in the sieve were removed from the substrate, placed individually into 1 oz cups, and covered with 2 g of their same substrate and not handled again. Non-handled 64 oz cups were removed from, and returned to, the incubator at the same time as handled cups but otherwise not touched.

To assess time to eclosion, each non-handled 64 oz cup, and all 1 oz cups of previously-handled pupae, were checked for any adult flies each day until day 30. Any adults found were thus 0-24 hours old at the time of collection. They were immediately removed from the cup, anesthetised with isoflurane, weighed to the nearest 0.1 mg (VWR-220TC), sexed, and the day of eclosion was recorded. Within 24 hours of collection, the ventral abdominal window of each fly was photographed against a red background under a dissecting microscope at 2X magnification (AmScope, SM-1TSZ-V203) in order to visualise the fat body and determine ‘window fullness’ (Figure 1; Harjoko *et al*., 2023; Oliveira *et al*., 2016). Flies were then frozen and stored at 4 °C.

**Figure 1.**
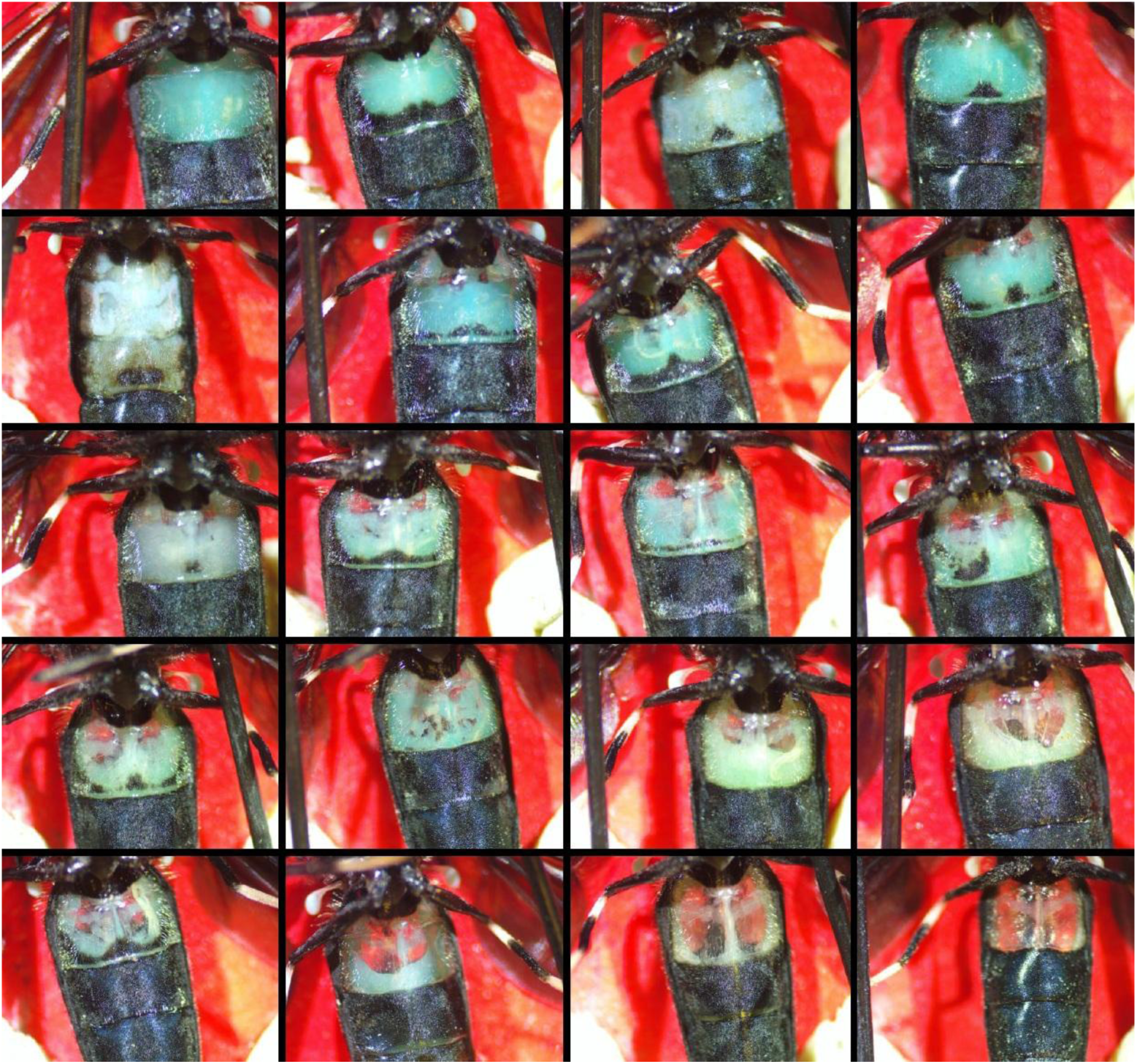
Examples of window fullness photos (2X magnification) from the ventral surface of the BSF abdomen. The fat body tissue is visualised as blue against the red background; entomological pins and modeling clay were used to hold the legs and wings out of the way for the photos.

Eclosion rate was calculated using the formula:

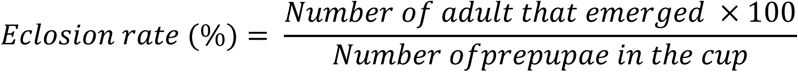

On day 30, we removed each handled pupae from its 1 oz cup and dissected it with microscissors. All individuals that had not yet eclosed, but had pupated, were dead. In addition, we went through each 64 oz cup of handled and non-handed and noted the condition of the remaining individuals in each cup (live/dead prepupae; live/dead pupae).

### Adult body size: head width, thorax length, and dry mass

The head width of the adult was measured as the distance between the widest parts of the head to the nearest 0.1 mm with a digital caliper (WEN). The thorax length was measured as the distance between the midpoint of the prescutum and the midpoint at the end of the scutellum to the nearest 0.1 mm. After these measurements were taken, flies were placed individually in open 1.5 mL plastic vials and put inside a convection oven at 60°C for 72 hr. Dry mass was then recorded to the nearest 0.1 mg on the analytical balance.

### Analyzing window fullness photos

Photos of fly windows were analyzed using ImageJ’s freehand selection tool (Figure 1; Schneider *et al*., 2012) to quantify the total window area and the fat body area. Relative window fullness was calculated as in Harjoko *et al*. (2023), but as a proportion, using the formula:

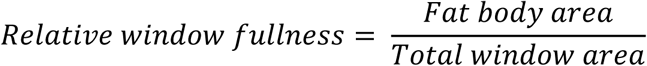

### Statistical Analyses

All analyses were performed using R (version 4.4.2) while charts were made in GraphPad Prism (version 10.2.3). Alpha was set to 0.05. All data were tested for normality and heteroskedasticity. We analyzed data for each sex independently, as BSF sexes are known to be dimorphic in development and morphology. Wet mass, dry mass, head width, and thorax length were analyzed using a generalised linear mixed model (GLMM) with a gamma distribution and log link function, (including substrate, moisture content, and handling) with cup as a random factor. Emergence day was analyzed using a GLMM with a poisson distribution and log link function (McCullagh, 1989), and using cup as a random factor.

Eclosion rate and window fullness were analyzed using beta regressions (preferred for data that are bounded at 0 and 1; Zeileis *et al*., 2016), with cup used as a random factor in the analysis of window fullness. For all models, we used a stepwise backwards elimination approach based on p-values (Chowdhury and Turin, 2020) to build the model of best fit for each variable, beginning with all main effects and two-way interactions and eliminating variables that did not improve the fit of the model, beginning with two-way interactions. When a main effect was not significant but was involved in a significant two-way interaction, it was retained in the model. A posthoc Tukey’s multiple comparisons test (MCT) was used to separate means. The Emtrends function was used to assess if a trendline was significantly different from 0.

### Ethical use of insects in research statement

Research on insects is not currently subject to any legally-mandated ethical review in the United States; therefore, no ethical approval was required to conduct this study. Nevertheless, we aimed to apply the 3Rs (replacement, reduction, and refinement) where possible. We calculated the appropriate sample size for reduction using a power analysis in G*Power (power = 0.8, alpha = 0.05). We also used isoflurane to anesthetise adults prior to handling, restraint, and euthanasia as a refinement.

## Results

### Development Time and Eclosion Rate

Only handling affected development time, e.g., eclosion day, in both male (GLMM, poisson with log link; n = 700, Χ^2^ = 98.11, p < 0.0001) and female (n = 664, Χ^2^ = 104.23, p < 0.0001) flies. Handling delayed eclosion by an average of 3.3 days in males (20.4 ± 0.27 handled vs. 17.1 ± 0.2 days non-handled) and 3.5 days in females (20.5 ± 0.28 handled vs. 17 ± 0.21 days non-handled; Figure 2).

**Figure 2.**
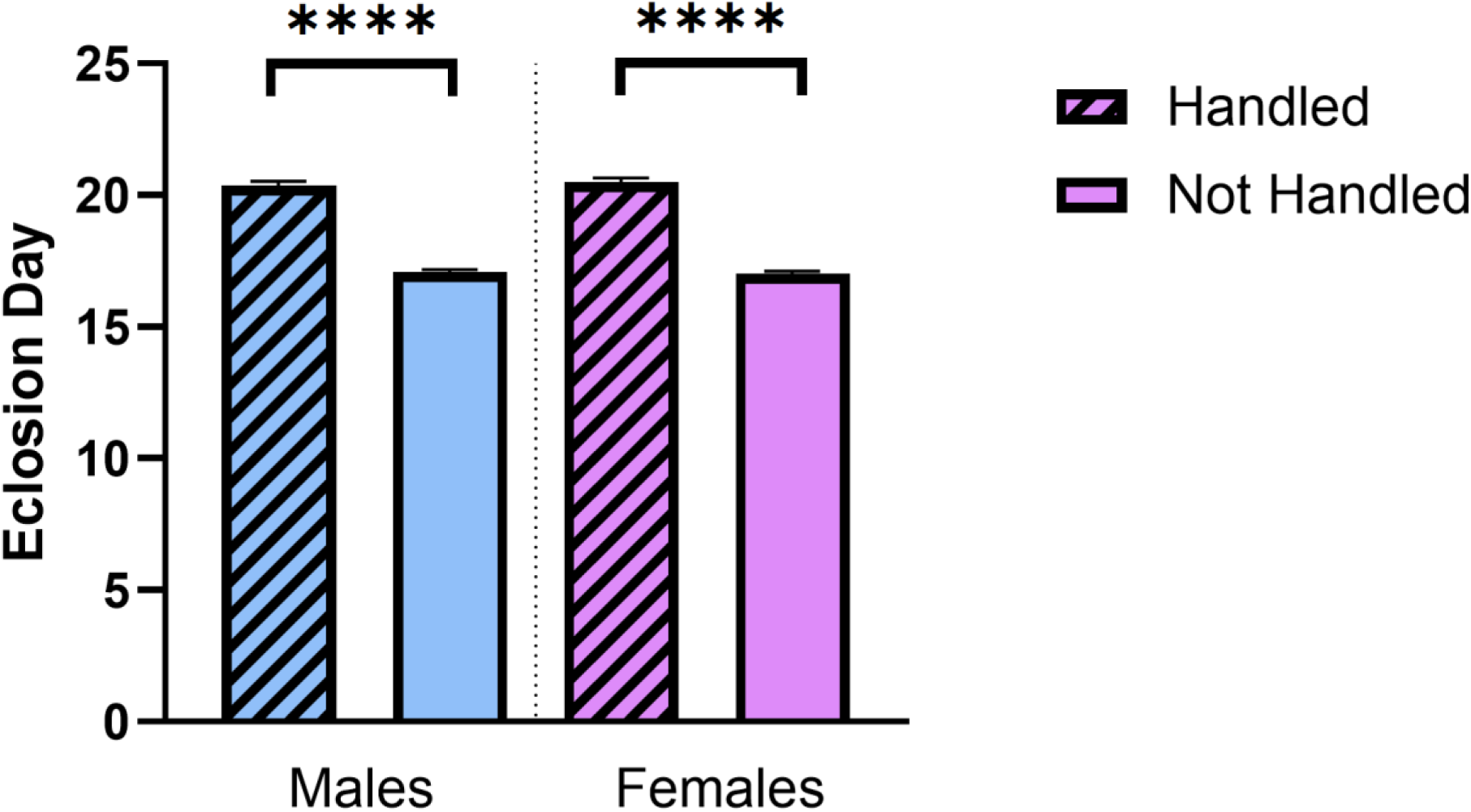
Eclosion day is delayed by > 3 days as a result of daily handling in BSF prepupae. Substrate, moisture content, and all two-way interactions were not significant predictors of eclosion day. (Males: GLMM, poisson with log link; n = 700, Χ^2^ = 98.11, p < 0.0001; females: n = 664, Χ^2^ = 104.23, p < 0.0001). **** = p < 0.0001. Bar graph shows mean and SE. Substrate (beta regression; n = 90; Χ^2^ = 10.81, p = 0.029), handling (X^2^ = 147.33, p < 0.0001), and the interactions of both substrate and handling (Χ^2^ = 66.94, p < 0.0001) and handling and moisture content (Χ^2^ = 9.09, p = 0.0026) significantly impacted eclosion rate (Table 1). FS had a significantly lower eclosion rate (72.1 ± 2.4%) than CC (82.2 ± 2.3%) and WC (81.5 ± 2.2%); PS and VM were intermediate (Tukey’s MCT; Table 1).

**Table 1.** Impacts of substrate, handling, interaction between substrate and handling, and interaction between handling and moisture content on eclosion rate in flies.

|  | **Eclosion rate (%)** |
| --- | --- |
| **Substrate\*** |  |
| Corn cob grits (CC) | $82.2 \pm 2.3^a$ |
| Frass (FS) | $72.1 \pm 2.4^b$ |
| Potting soil (PS) | $74.3 \pm 2.3^{ab}$ |
| Vermiculite (VM) | $75.9 \pm 2.6^{ab}$ |
| Wood chips (WC) | $81.5 \pm 2.2^a$ |
| **Handling\*** |  |
| Handled | $61.9 \pm 1.9^b$ |
| Non-handled | $92.5 \pm 0.9^a$ |
| **Substrate × handling\*** |  |
| CC × handled | $75.3 \pm 3.8^{cde}$ |
| FS × handled | $49.4 \pm 4.5^f$ |
| PS × handled | $50.0 \pm 4.5^f$ |
| VM × handled | $68.6 \pm 4.1^{def}$ |
| WC × handled | $66.4 \pm 4.2^{ef}$ |
| CC × non-handled | $89.2 \pm 2.5^{bc}$ |
| FS × non-handled | $94.9 \pm 1.5^{ab}$ |
| PS × non-handled | $98.6 \pm 0.5^a$ |
| VM × non-handled | $83.2 \pm 3.2^{cd}$ |
| WC × non-handled | $96.5 \pm 1.1^{ab}$ |
| **Handling × moisture\*\*** |  |
| Handled × overall moisture content | $-0.18 \pm 0.06$ ($p = 0.0023$) |
| Non-handled × overall moisture content |  |

\*Letters indicate statistically significant differences among groups, reporting model-adjusted mean and SE (Tukey's).
\*\*The trend (slope) was reported for an interaction that involved moisture content when significantly different ( $p < 0.05$ ) from zero.

Handling significantly reduced eclosion rate in every substrate except CC and VM (all p < 0.05; Figure 3), with an overall reduction in mean eclosion rate across all substrates of 30.6% (61.9 ± 1.9% in handled cups v. 92.5 ± 0.9% in non-handled cups). The impact of handling was greatest in PS (48.6% reduction in eclosion rate) and FS (45.5% reduction in eclosion rate). In the handled condition, an increase in moisture content decreased the proportion of BSF that eclosed (p = 0.0023; Table 1) while in non-handled conditions there was no effect of moisture content on the proportion of BSF that eclosed (p = 0.12).

**Figure 3.**
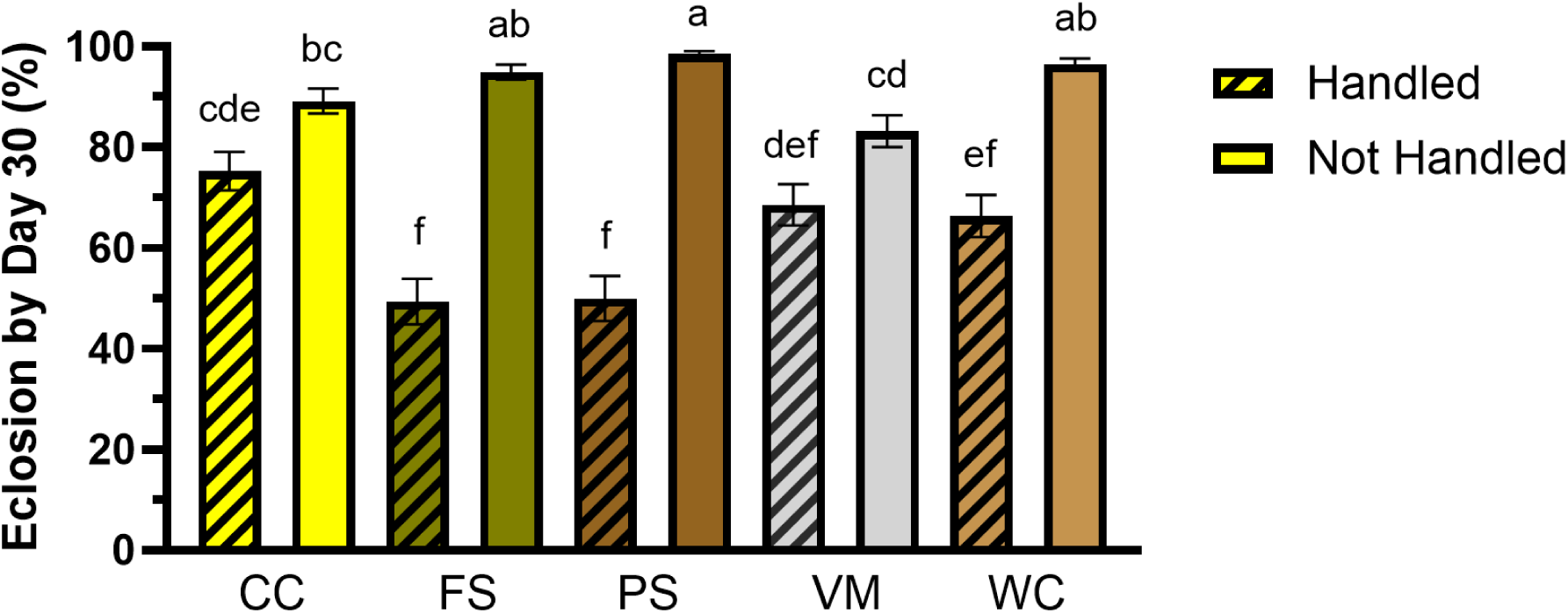
Substrate and handling affected the proportion of BSF adults that eclosed by day 30. Substrate type (beta regression; n = 90; Χ^2^ = 10.81, p = 0.029), handling (X^2^ = 147.33, p < 0.0001) and the interactions of substrate type and handling (Χ^2^ = 66.94, p < 0.0001) all affected eclosion by day 30. Handling significantly reduced eclosion rate in all substrates but CC and VM. Letters indicate statistically significant differences among groups (Tukey’s; p < 0.05). Bar graph shows model-adjusted means and SE.

Of the individuals that had not yet eclosed by the end of the study, the sample size was very small in most condition (at least 50% of conditions had less than 12 individuals for all three replicates combined, and some conditions had an n = 0). Therefore, we did not do statistical analyses. However, two observations were made from the raw data: 1) live prepupae were much more likely to be found on day 30 in the non-disturbed conditions than the disturbed conditions, where dead prepupae and, especially, dead pupae were common; and 2) that live prepupae were more likely to be found in vermiculite than other conditions, even when disturbed (Supplementary Table S2).

### Adult morphology: Body mass

Only substrate type affected adult female wet mass (n = 664, Χ^2^ = 28.63, p < 0.0001) and dry mass (n = 669, Χ^2^ = 11.64, p = 0.02; Table 2; Figure 4A, C). Females had the largest wet mass in CC (78.0 ± 1.3 mg) and WC (81.0 ± 1.4 mg), and the largest dry mass in WC (31.0 ± 0.6 mg).

**Figure 4.**
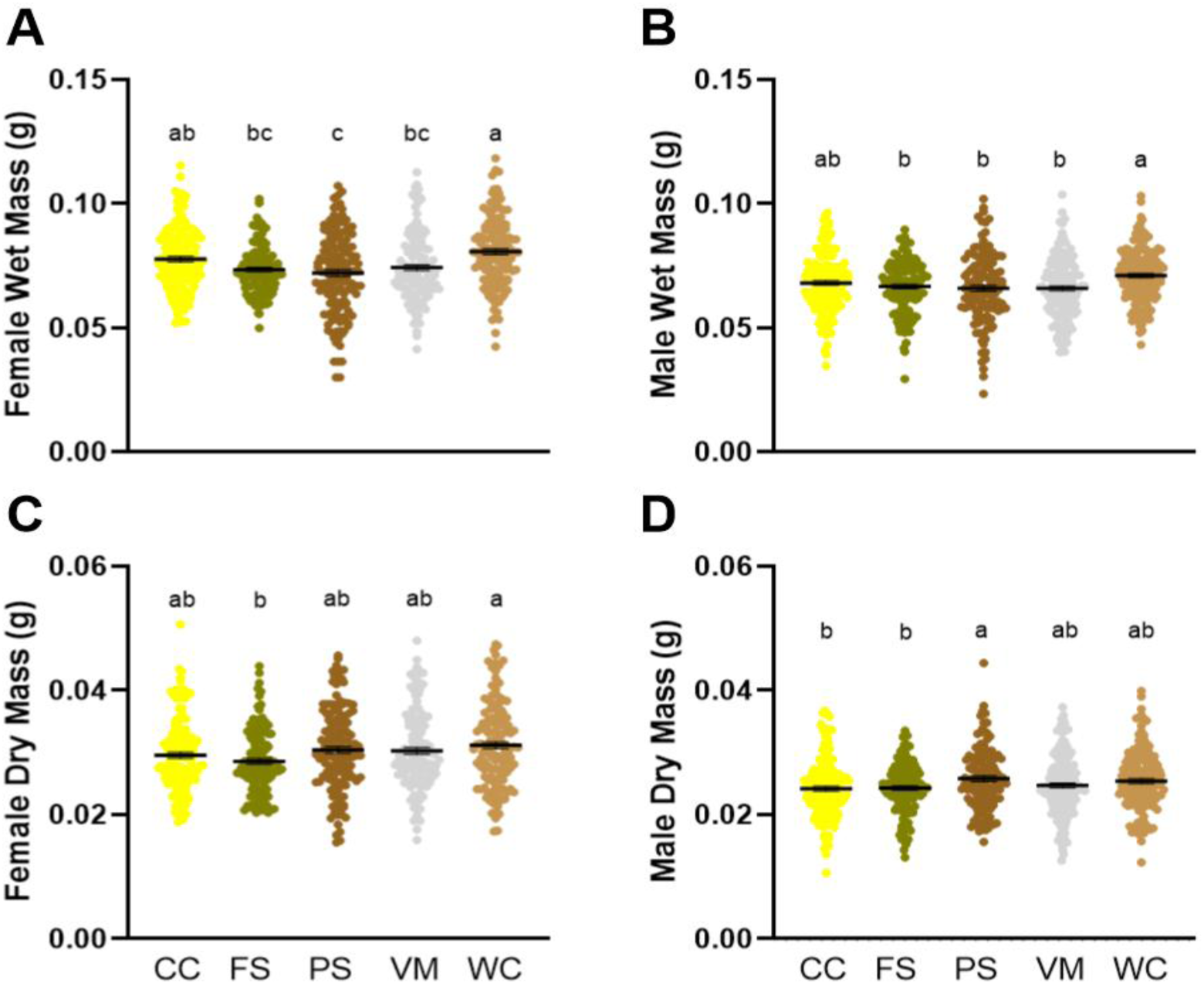
Substrate type significantly affected female (A,C) and male (B,D) mass. A, C) WC resulted in the highest mean wet and dry mass for females; FS and PS resulted in the lowest wet mass, and FS also resulted in reduced dry mass. B, D) WC resulted in the highest mean wet mass for males, while PS resulted in a lower wet mass but highest dry mass; FS also resulted in reduced wet and dry mass for males. Letters on graphs indicate statistically significant differences among conditions (Tukey’s; p < 0.05); horizontal black bar represents the mean with SE.

**Table 2.** Impacts of substrate, handling, interaction between substrate and handling, and interaction between substrate and moisture content on wet and dry mass in flies.

|  | Male wet mass | Female wet mass | Male dry mass | Female dry mass |
| --- | --- | --- | --- | --- |
| **Substrate\*** |  |  |  |  |
| CC | 0.068 ± 0.0010<sup>ab</sup> | 0.078 ± 0.0013<sup>ab</sup> | 0.024 ± 0.00037<sup>b</sup> | 0.030 ± 0.00054<sup>ab</sup> |
| FS | 0.066 ± 0.0011<sup>b</sup> | 0.073 ± 0.0014<sup>bc</sup> | 0.024 ± 0.00041<sup>b</sup> | 0.029 ± 0.00057<sup>b</sup> |
| PS | 0.065 ± 0.0011<sup>b</sup> | 0.072 ± 0.0012<sup>c</sup> | 0.026 ± 0.00049<sup>a</sup> | 0.030 ± 0.00056<sup>ab</sup> |
| VM | 0.066 ± 0.0010<sup>b</sup> | 0.074 ± 0.0013<sup>bc</sup> | 0.025 ± 0.00039<sup>ab</sup> | 0.030 ± 0.00057<sup>ab</sup> |
| WC | 0.071 ± 0.0010<sup>a</sup> | 0.081 ± 0.0014<sup>a</sup> | 0.025 ± 0.00039<sup>ab</sup> | 0.031 ± 0.00058<sup>a</sup> |
| **Handling\*** |  |  |  |  |
| Handled | 0.066 ± 0.00071<sup>b</sup> |  |  |  |
| Non-handled | 0.069 ± 0.00061<sup>a</sup> |  |  |  |
| **Substrate × handling\*** |  |  |  |  |
| CC × handled |  |  | 0.024 ± 0.00056<sup>ab</sup> |  |
| FS × handled |  |  | 0.024 ± 0.00065<sup>ab</sup> |  |
| PS × handled |  |  | 0.027 ± 0.00083<sup>a</sup> |  |
| VM × handled |  |  | 0.024 ± 0.00057<sup>b</sup> |  |
| WC × handled |  |  | 0.024 ± 0.00055<sup>b</sup> |  |
| CC × non-handled |  |  | 0.024 ± 0.00050<sup>b</sup> |  |
| Frass × non-handled |  |  | 0.024 ± 0.00052<sup>ab</sup> |  |
| PS × non-handled |  |  | 0.025 ± 0.00052<sup>ab</sup> |  |
| VM × non-handled |  |  | 0.025 ± 0.00054<sup>ab</sup> |  |
| WC × non-handled |  |  | 0.027 ± 0.00055<sup>a</sup> |  |
| **Substrate × moisture\*\*** |  |  |  |  |
| CC × moisture |  |  |  |  |
| FS × moisture |  |  |  |  |
| PS × moisture | 0.0016 ± 0.00048<br>(p = 0.0009) |  | 0.0012 ± 0.00050<br>(p = 0.02) |  |
| VM × moisture | -0.0013 ± 0.00045<br>(p = 0.0038) |  | -0.0016 ± 0.00048<br>(p = 0.0006) |  |
| WC × moisture |  |  |  |  |

\* Model-adjusted mean and SE (in g) reported.
\*\*The trend (slope) was reported for an interaction that involved moisture content when significantly different (p < 0.05) from zero.

Substrate (Χ^2^ = 19.77, p = 0.0004), handling (Χ^2^ = 11.87, p = 0.0006) and interaction between substrate and moisture content (Χ^2^ = 23.32, p = 0.0001) affected the wet mass of male flies (n = 700, Table 2; Figure 4B). Substrate (Χ^2^ = 11.36, p = 0.023), the interaction between substrate and moisture content (Χ^2^ = 18.97, p = 0.0008), and the interaction between substrate and handling (Χ^2^ = 11.6, p = 0.02) affected the dry mass of male flies (Table 2; Figure 4D).

Males had the largest wet mass in WC (71.0 ± 1.0 mg) and CC (68.0 ± 1.0 mg), and the largest dry mass in PS (26.0 ± 0.5 mg). FS significantly reduced wet (66.0 ± 1.1 mg) and dry (24.0 ± 0.4 mg) mass, and dry mass was also reduced in CC (24.0 ± 0.4 mg) in males. Handling reduced mean wet mass by 3.0 mg (4.3%) but had no effect on dry mass (p > 0.05) in males. For every 1% increase in moisture content, the wet mass of male flies that emerged from PS increased by 1.60 mg (p = 0.0009) and the dry mass increased by 1.2 mg (p = 0.02). Conversely, the wet mass of male flies that emerged from VM decreased by 1.3 mg (p = 0.0038) and the dry mass decreased by 1.6 mg (p = 0.0006). However, moisture content had no significant effect on the wet or dry mass of male flies that pupated in the other substrates (p > 0.05 for all).

### Adult morphology: Head width

Substrate (males: n = 700, Χ^2^ = 17.16, p = 0.0018; females: n = 664, Χ^2^ = 34.55, p < 0.0001), handling (males: Χ^2^ = 7.75, p = 0.0054; females: Χ^2^ = 4.46, p = 0.0347), the interaction of substrate and handling (males: Χ^2^ = 20.88, p = 0.0003; females: Χ^2^ = 15.11, p = 0.004), and the interaction of handling and moisture (males: Χ^2^ = 19.92, p < 0.0001; females: Χ^2^ = 11.35, p < 0.0001), affected head width in both sexes (Table 3).

**Table 3.** Impacts of substrate, handling, interaction between substrate and handling, and interaction between substrate and moisture content on head width in flies.

|  | Male head width | Female head width |
| --- | --- | --- |
| **Substrate\*** |  |  |
| CC | $3.65 \pm 0.02^{ab}$ | $3.81 \pm 0.03^b$ |
| FS | $3.59 \pm 0.03^b$ | $3.73 \pm 0.03^c$ |
| PS | $3.64 \pm 0.03^{ab}$ | $3.82 \pm 0.03^b$ |
| VM | $3.59 \pm 0.02^b$ | $3.73 \pm 0.03^c$ |
| WC | $3.71 \pm 0.03^a$ | $3.93 \pm 0.03^a$ |
| **Handling\*** |  |  |
| Handled | $3.60 \pm 0.02^b$ | $3.78 \pm 0.02^b$ |
| Non-handled | $3.67 \pm 0.01^a$ | $3.83 \pm 0.02^a$ |
| **Substrate × handling\*** |  |  |
| CC × handled | $3.67 \pm 0.04^{ab}$ | $3.83 \pm 0.04^{ab}$ |
| FS × handled | $3.50 \pm 0.04^c$ | $3.63 \pm 0.04^c$ |
| PS × handled | $3.64 \pm 0.04^{abc}$ | $3.79 \pm 0.04^{bc}$ |
| VM × handled | $3.58 \pm 0.03^{bcd}$ | $3.75 \pm 0.04^{bc}$ |
| WC × handled | $3.63 \pm 0.03^{bc}$ | $3.88 \pm 0.04^{abc}$ |
| CC × non-handled | $3.62 \pm 0.03^{bc}$ | $3.78 \pm 0.04^{bc}$ |
| FS × non-handled | $3.69 \pm 0.03^{ab}$ | $3.83 \pm 0.04^{ab}$ |
| PS × non-handled | $3.64 \pm 0.03^{bc}$ | $3.86 \pm 0.04^{ab}$ |
| VM × non-handled | $3.60 \pm 0.03^{bc}$ | $3.70 \pm 0.04^{bc}$ |
| WC × non-handled | $3.80 \pm 0.03^a$ | $3.99 \pm 0.04^a$ |
| **Substrate × moisture\*\*** |  |  |
| CC × moisture | | |
| FS × moisture | $-0.00044 \pm 0.00022$ ($p = 0.047$) | |
| PS × moisture | $0.00045 \pm 0.00021$ ($p = 0.036$) | |
| VM × moisture | | |
| WC × moisture | | |
| **Handling × moisture\*\*** |  |  |
| Handled × moisture | $0.00051 \pm 0.00014$ ($p = 0.0003$) | $0.00043 \pm 0.00015$ ($p = 0.004$) |
| Non-handled × moisture | $-0.00032 \pm 0.00012$ ($p = 0.009$) | |

\* Model-adjusted mean and SE (in mm) reported.
\*\*The trend (slope) was reported for an interaction that involved moisture content when significantly different ( $p < 0.05$ ) from zero.

The widest heads were observed in WC for both male (3.71 ± 0.03 mm) and female (3.93 ± 0.03 mm) flies. Head width was significantly reduced in FS and VM for both males (FS: 3.59 ± 0.03; VM: 3.59 ± 0.02 mm) and females (FS and VM, both: 3.73 ± 0.03 mm). Handling reduced mean head width by 0.07 mm (1.9%) in males and by 0.05 mm (1.3%) in females (Figure 5). As a result of these factors, head widths were highest in non-handled WC for males (3.80 ± 0.03 mm) and females (3.99 ± 0.04 mm) and lowest in handled FS (males 3.5 ± 0.04 mm; females: 3.63 ± 0.04 mm). In handled conditions, increasing moisture content increased the head widths of male (p = 0.0003) and female (p = 0.004) flies. When not handled, an increase in moisture content decreased the head width of male flies (p = 0.009), but had no effect on the head width of non-handled female flies (p = 0.07).

**Figure 5.**
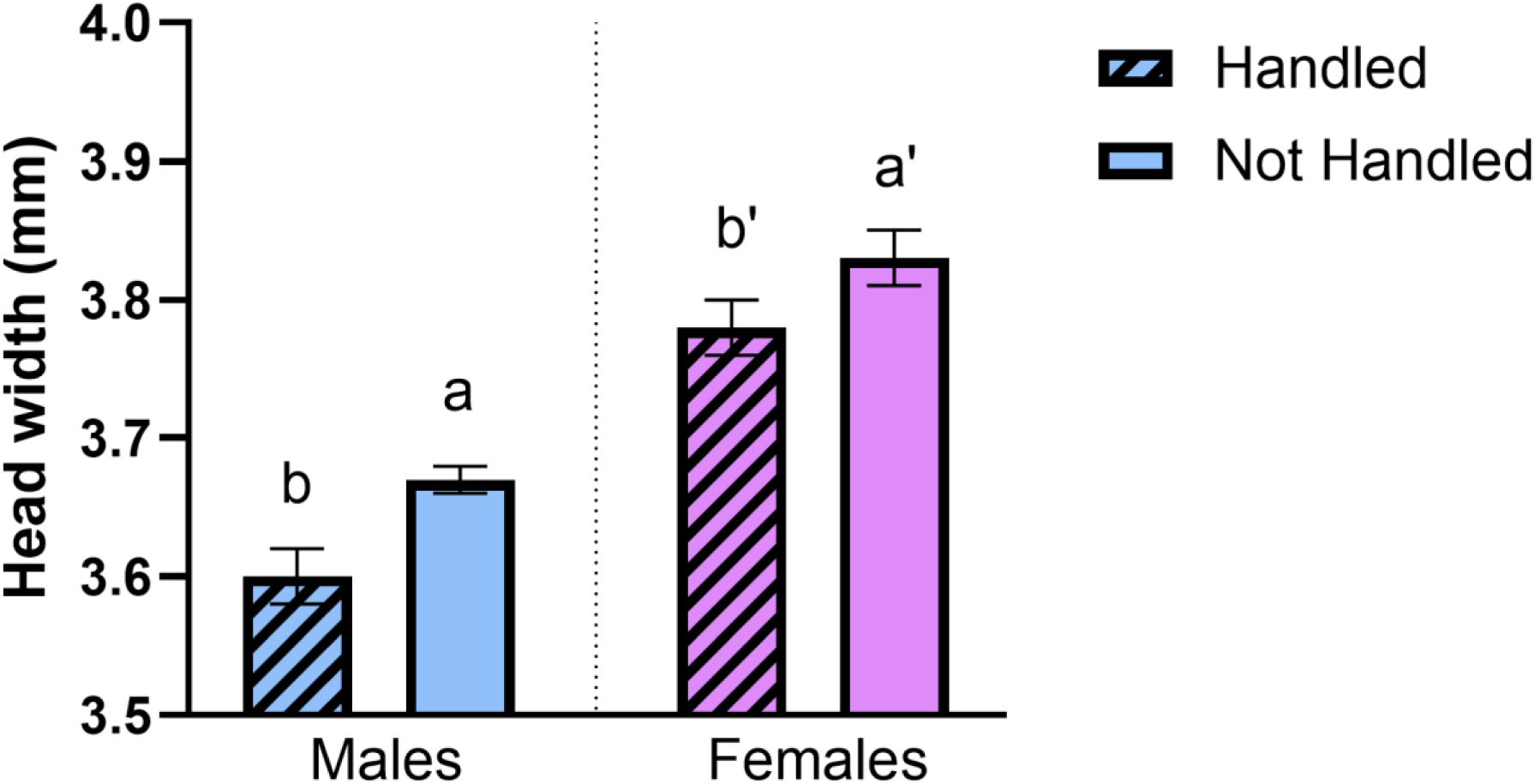
Handling reduced head width in both male (1.9%) and female (1.3%) flies. Letters on graphs indicate statistically significant differences among conditions (Tukey’s; p < 0.05); bar graph shows model-adjusted mean with SE.

Male head width was also affected by the interaction of substrate and moisture (Χ^2^ = 13.59, p = 0.0087). An increase in moisture content increased the head width of male flies in PS (p =0.036) and decreased head width of male flies in FS (p = 0.047).

### Adult morphology: Thorax length

Substrate affected thorax length in both male (n = 700, Χ^2^ = 39.35, p < 0.0001) and female (n = 663, X^2^ = 14.37, p = 0.0062) flies (Table 4). Thorax lengths were highest in PS and WC for both male (PS: 5.34 ± 0.04 mm; WC: 5.35 ± 0.04 mm) and female (PS: 5.14 ± 0.04; WC: 5.17 ± 0.04) flies.

**Table 4.** Impacts of substrate, interaction between substrate and moisture content, and interaction between handling and moisture content on thorax length in flies.

|  | Male thorax length | Female thorax length |
| --- | --- | --- |
| **Substrate\*** |  |  |
| CC | 5.15 ± 0.04<sup>b</sup> | 5.04 ± 0.04<sup>ab</sup> |
| FS | 5.19 ± 0.04<sup>ab</sup> | 5.05 ± 0.04<sup>ab</sup> |
| PS | 5.34 ± 0.04<sup>a</sup> | 5.14 ± 0.04<sup>a</sup> |
| VM | 5.06 ± 0.04<sup>b</sup> | 4.98 ± 0.04<sup>b</sup> |
| WC | 5.35 ± 0.04<sup>a</sup> | 5.17 ± 0.04<sup>a</sup> |
| **Substrate × moisture\*\*** |  |  |
| CC × moisture | -0.00054 ± 0.00023 (p = 0.018) |  |
| FS × moisture |  |  |
| PS × moisture |  |  |
| VM × moisture | -0.00052 ± 0.00024 (p = 0.03) |  |
| WC × moisture |  |  |
| **Handling × moisture\*\*** |  |  |
| Handled × moisture |  |  |
| Non-handled × moisture | -0.00037 ± 0.00014 (p = 0.011) |  |

\*Model-adjusted mean and SE (in mm) reported.
\*\*The trend (slope) was reported for an interaction that involved moisture content when significantly different (p < 0.05) from zero.

The interaction between substrate and moisture (Χ^2^ = 16.45, p = 0.0025) and the interaction between moisture and handling (Χ^2^ = 8.83, p = 0.003) also affected thorax length in male flies. The thorax length of males flies in CC (p = 0.018) and VM (p = 0.03) decreased as moisture content increased. In non-handled conditions, increasing moisture content decreased male thorax length (but had no effect in handled conditions; Table 4).

### Window Fullness

The interaction of substrate and handling affected window fullness in male (Χ^2^ = 21.19, p = 0.0003) and female (Χ^2^ = 22.55, p = 0.0002) flies (Table 5). Male windows were relatively more full of fat body in non-handled FS (0.72 ± 0.03) while female windows were most full in non-handled CC (0.94 ± 0.008). Across both sexes, reduced window fullness was observed in CC (handled) and PS (not handled).

**Table 5.**
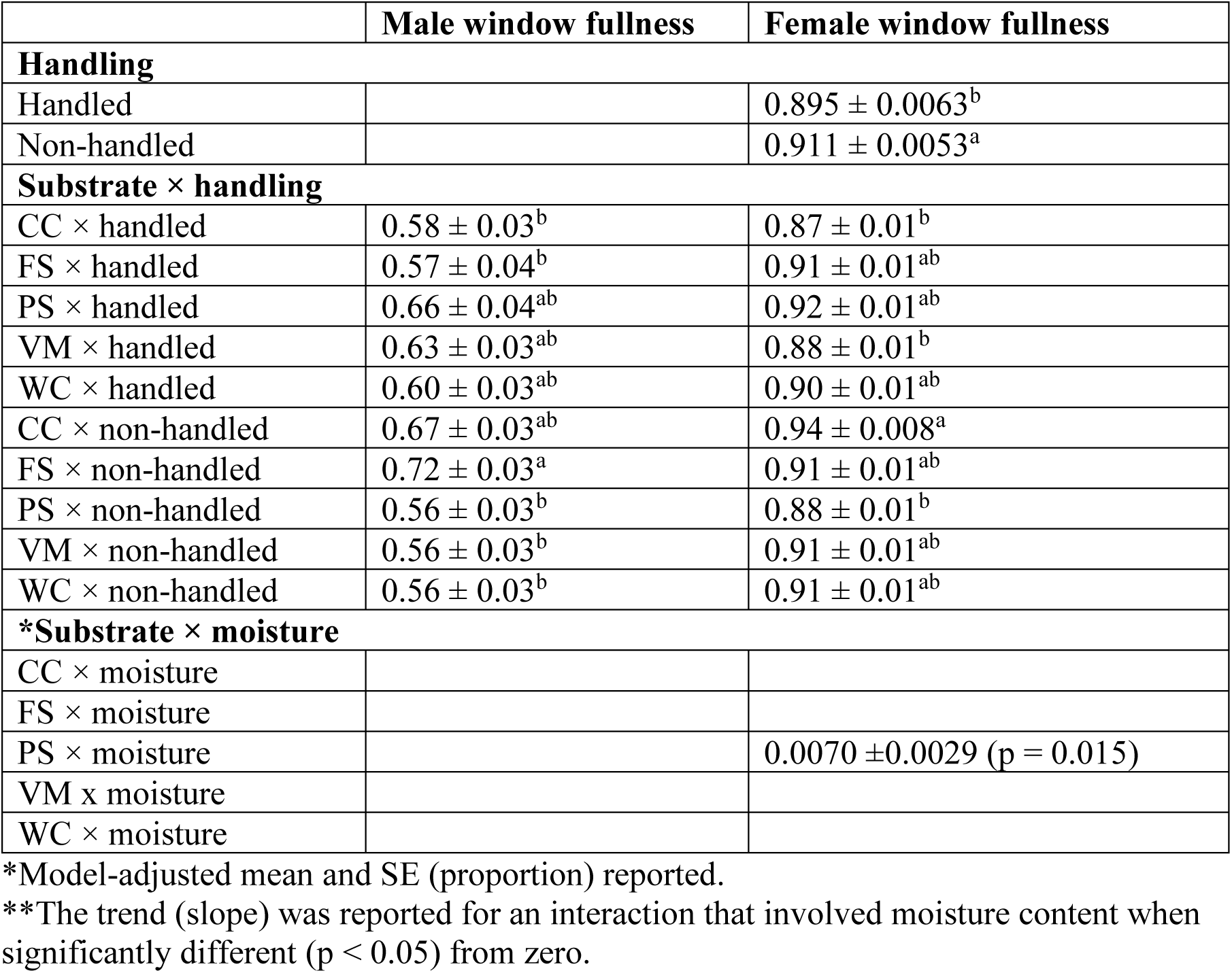
Impacts of handling, interaction between substrate and handling, and interaction between substrate and moisture content on window fullness in flies.

|  | Male window fullness | Female window fullness |
| --- | --- | --- |
| **Handling** |  |  |
| Handled | | $0.895 \pm 0.0063^b$ |
| Non-handled | | $0.911 \pm 0.0053^a$ |
| **Substrate × handling** |  |  |
| CC × handled | $0.58 \pm 0.03^b$ | $0.87 \pm 0.01^b$ |
| FS × handled | $0.57 \pm 0.04^b$ | $0.91 \pm 0.01^{ab}$ |
| PS × handled | $0.66 \pm 0.04^{ab}$ | $0.92 \pm 0.01^{ab}$ |
| VM × handled | $0.63 \pm 0.03^{ab}$ | $0.88 \pm 0.01^b$ |
| WC × handled | $0.60 \pm 0.03^{ab}$ | $0.90 \pm 0.01^{ab}$ |
| CC × non-handled | $0.67 \pm 0.03^{ab}$ | $0.94 \pm 0.008^a$ |
| FS × non-handled | $0.72 \pm 0.03^a$ | $0.91 \pm 0.01^{ab}$ |
| PS × non-handled | $0.56 \pm 0.03^b$ | $0.88 \pm 0.01^b$ |
| VM × non-handled | $0.56 \pm 0.03^b$ | $0.91 \pm 0.01^{ab}$ |
| WC × non-handled | $0.56 \pm 0.03^b$ | $0.91 \pm 0.01^{ab}$ |
| **\*Substrate × moisture** |  |  |
| CC × moisture | | |
| FS × moisture | | |
| PS × moisture | | $0.0070 \pm 0.0029$ ($p = 0.015$) |
| VM × moisture | | |
| WC × moisture | | |

\*Model-adjusted mean and SE (proportion) reported.
\*\*The trend (slope) was reported for an interaction that involved moisture content when significantly different ( $p < 0.05$ ) from zero.

Handling (Χ^2^ = 6.15, p = 0.013) and the interaction between substrate and moisture (Χ^2^ = 10.37, p = 0.035) also significantly affected window fullness in female flies. Handling reduced (p = 0.029) window fullness in female flies by 1.8% (Table 5). In PS only, an increase in moisture content also increased female window fullness (p = 0.015).

## Discussion

Our results suggest that prepupal handling and pupation substrate type have significant impacts on development time and subsequent adult morphology in BSF. Short bouts of daily handling during the prepupal phase negatively affected survival, development and body size: handling reduced eclosion rate by 30.6%, increased development time in both sexes by more than 3 days, decreased male wet mass by 4.3%, decreased head widths in both males (1.9%) and females (1.3%), and decreased window fullness in females by 1.8%. The largest effects of handling on eclosion rate were observed in FS and PS. Our results mirror data from other dipterans: Kobayashi and Takanashi (2025) also found that vibrations suppressed larval development and reduced adult emergence in dark-winged fungus gnats (Diptera: Sciaridae).

The strong effects of handling on eclosion rates and adult morphology may result from a number of factors, particularly stress and/or damage. Predator-linked cues, like vibrations and/or physical touch, are known to cause significant stress in insects (Cinel *et al*., 2020). Acute stress can lead to the activation of the octopamine-adipokinetic hormone axis (e.g., Adamo and Baker, 2011), resulting in increased metabolic rates and thereby energy consumption as the insect attempts to remove itself from the acute stressor. Octopamine has been tentatively linked to handling in BSF larvae (unpublished work: Baumann, 2019) and is known to cause mobilization of lipid reserves, like fat body tissue, in many species of insects (Arrese and Soulages, 2010). Additionally, handling can activate the cellular immune response: Tokusumi *et al*. (2018) reported stimulation of cellular immune response when *Drosophila melanogaster* (Diptera; Drosophilidae) larvae were squeezed with forceps. Immune activation is also energetically costly and can divert resources away from developmental processes (Adamo, 2014; Adamo, 2017a, b; Karpova *et al*., 2024; Zhang *et al*., 2019).

As prepupae are post-feeding, they cannot compensate for any increased energy burn during their prolonged development by consuming additional nutrients. Smaller energy reserves heading into pupation may explain our data demonstrating both smaller adults (for those that survive to eclosion) and a significant proportion of individuals (30.6%) not surviving the energy-intensive process of pupation at all. Somewhat in line with this hypothesis, the window fullness of newly eclosed female BSF, which can serve as an indicator of the nutritional status and energy/lipid resources of BSF (Harjoko *et al*., 2023), was reduced by 1.8% in handled conditions compared to non-handled conditions; coupled with a reduction in body size, this suggests smaller energy reserves for adults emerging from handled conditions. Window fullness may be a coarse but non-invasive proxy for the amount of stored metabolic resources in adult BSF.

The lower eclosion rates observed in handled flies could also have resulted from damage to the cuticle or internal structures during daily handling. Importantly, however, lower eclosion rates in handled conditions were largely due to prepupae not surviving pupation (at which point they were no longer being handled), not animals dying at the prepupal stage. Dead pupae that were dissected from handled conditions were almost all partially developed into adults, indicating the beginning of metamorphosis had occurred but that it could not be completed. Further, we rarely observed damage to the pupal case or the insect upon dissection of dead individuals that had pupated - though even small amounts of damage to the pupal case can increase the likelihood of disease or desiccation (Clark and Triblehorn, 2014; Kavaliers and Choleris, 2001; Parle *et al*., 2017; Sharp and Little, 1982; Taylor, 2020; Vincent and Wegst, 2004). Further, excessive vibration could disrupt airflow or damage tracheal tubes, chordotonal organs, imaginal discs, and other internal tissues (Nyberg and Muto, 2020; Tokusumi *et al*., 2018); the effects of this damage, if severe, could preclude subsequent normal eclosion even if metamorphosis begins. Finally, handling of the prepupae could result in oxidative stress and the increased production of reactive oxygen species that disrupt cellular, organ, or whole-body functions, leading to death (Kodrik *et al*., 2015).

Interestingly, prior research on BSF larvae found that daily handling either reduced development time (Nguyen *et al*., 2013) or did not affect it (Loiotine *et al*., 2024). However, our results demonstrate that daily handling at the prepupal stage prolonged development. These life-stage specific responses to predator-linked stressors might be adaptive: for instance, larvae that are exposed to a stressor may want to leave the stressful feeding substrate for a safer pupation area as rapidly as possible, whereas prepupae in a stressful environment may want to delay pupation and continue searching for a safer environment to pupate in. Irrespective of functional value, the variance in response to stress demonstrates how unique the behaviours of post-feeding prepupae and feeding larvae can be and justifies further research aimed at understanding the prepupal stage for both fundamental and applied work. Mechanistically, the prolonged adult emergence observed in handled flies might have resulted from the stress-induced shifts in hormone titres, especially for juvenile hormone (JH) and ecdysone (20E) that initiate and regulate metamorphosis (Chernysh, 2018; Perić-Mataruga *et al*., 2006; Smykal *et al*., 2014).

We also found that substrate type (given the five substrates we provided: corn cob grits, frass, potting soil, vermiculite, and wood chips) had significant impacts on BSF development and morphology, in line with prior research (Dzepe *et al*., 2020; Holmes *et al*., 2013). In our study, “Sani-chips” wood chips (WC), consistently performed above the other substrates with the highest eclosion rate (especially when not handled), highest wet mass, thorax length, and head widths for both males and females (and a high dry mass) and no significant difference from other substrates in window fullness. Sanichips are small wood chips used for laboratory rodent bedding and have few sharp edges; they thus have many characteristics of a desirable pupation substrate for BSF: loose enough to allow movement of prepupae, low compaction, non-toxic, and capable of retaining moisture (Pascacio-Villafán *et al*., 2021). Dzepe *et al*. (2020) also observed the highest emergence rate in wood shavings when comparing emergence success in wood shavings, fine sand, wheat bran, and no substrate. Holmes *et al*. (2013) found no difference in the adult emergence from sand, topsoil, wood shavings and potting soil. Wood chips/shavings have a variety of textures and moisture retention abilities; larger shavings with sharper edges are not preferred by BSF prepupae in two-choice assays (Durosaro, pers. obs.); the exact characteristics of the wood shavings used may thus explain discrepancies between studies.

Prior studies found that moisture content can affect pupation and adult emergence, with low moisture content (0 - 20%) resulting in increased mortality (Liu *et al*., 2023). Surprisingly, we did not observe an effect of low moisture content (20%) on any of our response variables, though moisture content did sometimes interact with substrate or handling type (with few consistent trends emerging). This may have been a result of the higher relative humidity in our incubator (∼71% RH) compared to other studies (∼50% RH), which could have supplemented the poor moisture content in the substrates themselves.

By contrast, BSF frass (FS) performed the worst with the lowest eclosion rate, lowest wet and dry mass, and lowest head width for both sexes. The poor performance of FS is perhaps unsurprising given that BSF prepupae naturally leave their own frass and seek out other substrates in which to pupate (Holmes *et al*., 2013). In some insects, like the yellow mealworm beetle (*Tenebrio molitor;* Coleoptera: Tenebrionidae), frass is known to contain compounds (JH) that actively suppress or delay pupation upon consumption, delaying pupation at high densities (Weaver and McFarlane, 1990). In BSF prepupae, however, FS did not delay development compared to the other substrates; further, the prepupae we tested were exclusively post-feeding and therefore could not have been consuming JH if it were found in the frass. Thus, the mechanism by which FS reduces eclosion rates and alters adult morphology when prepupae are exposed could be further researched.

Our data demonstrate that both prepupal handling and pupation substrate type can significantly affect the survival, development, and morphology of adult BSF. Practically, frequent bouts of handling are likely to be more common in laboratory settings than production settings (though handling events may be harsher in production settings with automated sieves). Our data thus highlight a gap between lab and production practices that could result in difficulty translating benchtop research to production scales, given that handling can result in phenotypic changes in adults. Our data suggest that prepupal handling can be stressful and reducing prepupal handling in either frequency or duration may improve prepupal welfare while also serving production and research goals (increased survival, faster development, larger adult males and females with more energy reserves). Dependent on cost and availability, producers may want to use small, rounded wood chips as a pupation substrate given their clear benefits for survival and adult morphology. The best eclosion rate for handled prepupae was observed in corn cob grits, which also scored intermediate on many morphological variables; therefore, producers that are live-shipping prepupae (a situation that may induce stress similar to that seen in our study) may want to use corn cob grits as the substrate as it may increase the animals’ resilience to mechanical stress. Frass as a pupation substrate, and the lack of a pupation substrate entirely (Holmes *et al*., 2013; Durosaro and Barrett, pers. comm.), should be avoided as they resulted in the poorest animal welfare and production-relevant outcomes in our work or prior literature.

## Conflicts of Interest

None

## Funding Statement

EA was supported as part of the First Year Science Apprenticeship Program at Indiana University Indianapolis.

## Acknowledgements

We thank Charlie Schmidt and Sofia Goodpaster for their assistance with data input and/or adult morphology measurements. We thank Elijah Persson-Gordon for assisting with setting up the experiment and some prepupal handling. We thank Craig Perl for the advice on statistical analysis and helping with the R code for beta regressions.

## Supplementary Materials

**Supplementary Table S1.** Dry-rehydrated moisture content, weight of substrate and amount of water added to the substrates.

| Substrate | dry-rehydrated moisture content (%) | Weight equivalent to 8 cm (g) | Equivalent amount of water added at different moisture contents (mL) |  |  |
| --- | --- | --- | --- | --- | --- |
|  |  |  | 20% | 60% | 100% |
| Corn cob grits | 1.18 | 230.00 | 42.8 | 133.7 | 224.6 |
| Potting soil | 3.94 | 84.00 | 13.0 | 45.2 | 77.5 |
| Vermiculite | 1.26 | 93.00 | 17.2 | 53.9 | 90.7 |
| Wood chips | 2.12 | 87.00 | 15.2 | 49.3 | 83.3 |
| Frass | 2.64 | 236.00 | 39.9 | 131.8 | 223.7 |

**Supplementary Table S2.** Proportion of BSF that had not eclosed by day 30 that were found as live prepupae (LPP), live pupae (LP), dead prepupae (DPP), dead pupae (DP), or dead adults still in the substrate (DA)

| Substrate | Moisture | Handled (n = ) | Pr(LPP) | Pr(LP) | Pr(DPP) | Pr(DP) | Pr(DA) |
| --- | --- | --- | --- | --- | --- | --- | --- |
| CC | 20 | H (n = 21) | 0.05 | 0 | 0.19 | 0.76 | 0 |
| CC | 60 | H (n = 17) | 0.06 | 0 | 0.06 | 0.88 | 0 |
| CC | 100 | H (n = 13) | 0 | 0 | 0 | 1 | 0 |
| FS | 20 | H (n = 21) | 0.05 | 0 | 0.19 | 0.76 | 0 |
| FS | 60 | H (n = 26) | 0.08 | 0 | 0.31 | 0.62 | 0 |
| FS | 100 | H (n = 39) | 0.05 | 0 | 0.08 | 0.87 | 0 |
| PS | 20 | H (n = 19) | 0 | 0 | 0.05 | 0.95 | 0 |
| PS | 60 | H (n = 29) | 0 | 0 | 0 | 1 | 0 |
| PS | 100 | H (n = 39) | 0 | 0 | 0.5 | 0.95 | 0 |
| VM | 20 | H (n = 15) | 0.07 | 0 | 0 | 0.93 | 0 |
| VM | 60 | H (n = 13) | 0.62 | 0 | 0 | 0.38 | 0 |
| VM | 100 | H (n = 18) | 0.5 | 0 | 0.19 | 0.5 | 0 |
| WC | 20 | H (n = 12) | 0.42 | 0 | 0 | 0.58 | 0 |
| WC | 60 | H (n = 20) | 0 | 0 | 0 | 1 | 0 |
| WC | 100 | H (n = 21) | 0.1 | 0 | 0.05 | 0.86 | 0 |
| CC | 20 | NH (n = 5) | 0.2 | 0 | 0 | 0.8 | 0 |
| CC | 60 | NH (n = 3) | 0.33 | 0 | 0 | 0.33 | 0.33 |
| CC | 100 | NH (n = 8) | 0 | 0 | 0.13 | 0.5 | 0.37 |
| FS | 20 | NH (n = 7) | 0.29 | 0 | 0.14 | 0.57 | 0 |
| FS | 60 | NH (n = 13) | 0.15 | 0 | 0.15 | 0.69 | 0 |
| FS | 100 | NH (n = 0) | n.d.* | n.d. | n.d. | n.d. | n.d. |
| PS | 20 | NH (n = 0) | n.d. | n.d. | n.d. | n.d. | n.d. |
| PS | 60 | NH (n = 2) | 0.5 | 0 | 0 | 0.5 | 0 |
| PS | 100 | NH (n = 0) | n.d. | n.d. | n.d. | n.d. | n.d. |
| VM | 20 | NH (n = 9) | 1 | 0 | 0 | 0 | 0 |
| VM | 60 | NH (n = 11) | 0.64 | 0.18 | 0 | 0.18 | 0 |
| VM | 100 | NH (n = 9) | 1 | 0 | 0 | 0 | 0 |
| WC | 20 | NH (n = 3) | 0.67 | 0.33 | 0 | 0 | 0 |
| WC | 60 | NH (n = 4) | 0.75 | 0 | 0 | 0.25 | 0 |
| WC | 100 | NH (n = 3) | 1 | 0 | 0 | 0 | 0 |

\*n.d. = no data, as there were no individuals that had not eclosed after day 30.

## Notes

### Competing Interest Statement

The authors have declared no competing interest.

